# Constitutively active STAT5b feminizes mouse liver gene expression

**DOI:** 10.1101/2022.02.14.480424

**Authors:** Dana Lau-Corona, Hong Ma, Cameron Vergato, Andre Sarmento-Cabral, Mercedes del Rio-Moreno, Rhonda D Kineman, David J Waxman

## Abstract

STAT5 is an essential transcriptional regulator of the sex-biased actions of growth hormone (GH) in the liver. Delivery of constitutively active STAT5 (STAT5_CA_) to male mouse liver using an engineered adeno-associated virus with high tropism for the liver is shown to induce widespread feminization of the liver, with extensive induction of female-biased genes and repression of male-biased genes, largely mimicking results obtained when male mice are given GH as a continuous infusion. Many of the gene expression changes observed were associated with STAT5 binding to liver chromatin, supporting the proposed direct role of persistently active STAT5 in continuous GH-induced liver feminization. The feminizing effects of STAT5_CA_ were dose-dependent; moreover, at higher levels, overexpression of STAT5_CA_ resulted in some histopathology not seen in continuous GH-infused male liver, including hepatocyte hyperplasia and distorted liver architecture. These findings establish that the persistent activation of STAT5 by GH that characterizes female liver is by itself sufficient to account for the female-biased expression of a majority of female-biased genes. Moreover, histological changes seen when STAT5_CA_ is overexpressed highlight the importance of carefully evaluating such effects before considering such STAT5 derivatives for therapeutic use in treating liver disease.

## Introduction

Sex-differences in pituitary growth hormone (GH) secretory patterns control the sex-biased expression of hundreds of genes in mouse liver (1) and contribute to the sexual dimorphism of liver physiology and pathology (2,3). In males, GH is secreted as a series of intermittent pulses followed by extended periods when circulating GH is undetectable, whereas in females, GH secretion is near continuous (1,4). Sex differences in liver gene expression are largely abolished when pituitary GH secretion is ablated by hypophysectomy, and can be substantially restored when exogenous GH is administered as a series of pulses (male circulating GH pattern), which confers a male pattern of expression, or as a continuous infusion (female GH pattern), which results in a female pattern of expression (5,6).

GH binding to its cell surface receptor activates the receptor-bound tyrosine kinase JAK2, which in turn phosphorylates and thereby activates the latent cytoplasmic transcription factor STAT5 (7,8). GH receptor/JAK2-activated STAT5 dimerizes and translocates to the nucleus, where it binds to thousands of sites in liver chromatin and induces transcription of many target genes (9,10). STAT5b, the major liver STAT5 form, plays a crucial role in liver metabolism and is an essential transcriptional regulator of the sex-dependent actions of GH in the liver (11,12). In male liver, repeated pulses of hepatic STAT5 activity, nuclear localization and transcriptional activity are observed and closely track the successive pulses of male pituitary gland GH release (13–15), whereas in female liver, STAT5 is persistently activated by the near-continuous female plasma GH profile (9,16). In whole body STAT5b knockout male mice, 90% of hepatic male-biased genes are repressed and 60% of hepatic female-biased genes are de-repressed (i.e., induced) (11). Further, STAT5 binds in a sex-biased manner to several thousand sites in mouse liver chromatin, with the set of male-biased STAT5 binding sites enriched nearby genes showing male-biased expression and the set of female-biased binding sites enriched nearby female-biased genes (9,17). Recent studies in a hepatocyte-specific STAT5a/STAT5b-KO mouse model (18) confirm these findings (19), supporting the proposal that the loss of hepatic STAT5, rather than an indirect feedback response to the loss of STAT5 in the hypothalamus or the pituitary gland, is responsible for the loss of liver sexual dimorphism seen in global STAT5b knockout mice.

Continuous infusion of GH in male mice (cGH treatment) substantially feminizes liver gene expression (20,21), most likely by a combination of two mechanisms. First, cGH overrides the endogenous male pulsatile plasma GH profile. This leads to widespread repression of male-biased genes that are positively regulated by male plasma GH pulses (class I male-biased genes) and to de-repression of female-biased genes that are negatively regulated by male GH pulses (class II female-biased genes) (22). Second, cGH treatment imposes a persistent, female-like pattern of plasma GH stimulation on male liver, which leads to the induction of female-biased genes that are positively regulated by the endogenous female GH pattern (class I female-biased genes) and represses male-biased genes that are negatively regulated by the female GH pattern (class II male-biased genes) (15,22). GH-activated STAT5 is a strong transcriptional activator of its well established direct target genes, including *Igf1* and *Socs2* (10,23), and similarly, can directly activate at least some class I male-biased genes when it is activated by a plasma GH pulse in male mouse liver (15). However, it is uncertain whether STAT5 signaling directly regulates other classes of sex-biased genes, most notably class I female-biased genes, which require a female plasma GH pattern for expression, or alternatively, whether other GH-stimulated but STAT5-independent signaling pathways downstream of GH receptor and JAK2 (24–26) mediate their sex-dependent expression. The requirement of STAT5 for sex-biased hepatic expression of both class I and class II sex-biased genes (21) does not entirely resolve this question, given the apparent perturbation of feedback mechanisms leading to changes in circulating GH patterns seen in both STAT5 knockout and GH receptor-deficient mouse models (18,27).

Here, we address these questions using AAV8-STAT5_CA_, a novel adeno-associated virus (AAV) vector that we engineered to deliver STAT5b with a point mutation in its SH2 domain (STAT5b Asn-642 to His) (28). This mutation renders the STAT5 protein constitutively active (‘CA’) and thus able to support robust expression of hepatic *Igf1* in the absence of GH (29). AAV8-STAT5_CA_ is based on AAV serotype 8, which has high specificity for infection of hepatocytes (30,31). Moreover, the STAT5b mutant is expressed from the hepatocyte-specific thyroxin-binding globulin promoter, which confers additional specificity for persistent expression of the constitutively active STAT5 in hepatocytes. We hypothesized that the persistent expression of active STAT5 protein in male mouse liver mimics the persistent activation of liver STAT5 seen in female liver and will feminize gene expression. Our findings reveal that the delivery of STAT5_CA_ to male mouse liver leads to widespread induction of female-biased genes and repression of male-biased genes, and thus feminization of the liver by persistent plasma GH stimulation can largely be attributed to the persistent activation of liver STAT5b.

## Materials and Methods

### Preparation of AAV8-STAT5_CA_

Wild-type mouse STAT5b expression plasmid with an N-terminal Flag tag was obtained from SinoBiological, Wayne PA (cat. # MG51116-NF). A point mutation at base 1963 (A to C), which changes Asn642 to His, was introduced into the STAT5b coding sequence by Genscript (Piscataway, NJ). This mutation confers constitutive activity (CA) to STAT5b, including the ability to induce *Igf1* gene expression in livers of hypophysectomized (GH-deficient) rats (28,29). The mutated plasmid was sent to Penn Vector Core (University of Pennsylvania, Philadelphia, PA), where the N-Flag-mSTAT5b_CA_ cDNA was cloned into the Penn Vector Core plasmid pENN.AAV.TBG.PI.eGFP.WPRE.bGH, replacing eGFP by the mutated and tagged STAT5b_CA_. The resultant plasmid, pENN.AAV.TBG.PI.N-FLAG-mSTAT5bCA.WPRE.bGH, was sequence verified then packaged in AAV8 virus, which was purified, quantified with respect to genome copy (GC) number, and supplied by Penn Vector as a ready to use vector, AAV8-TBGp-mStat5b_CA_, referred to here as AAV8-STAT5_CA_. The plasmid pENN.AAV.TBG.PI.N-FLAG-mSTAT5bCA.WPRE.bGH, required to prepare AAV8-STAT5_CA_ (see lab methods, below), is being deposited and will be made available through Addgene (https://www.addgene.org/).

### AAV8-STAT5_CA_ expression in mouse liver

Initial studies to validate AAV8-STAT5_CA_ were performed in male C57Bl/6 mice on a GHR^fl/fl^ background at Jesse Brown VA Medical Center (Chicago, IL) with Institutional Animal Care and Use Committee approval (protocol number 19-05). AAV8-TBGp-Null vector (Penn Vector Core), referred to here as AAV8-Null, was used as a control. Male GHR^fl/fl^ mice (10-12 wk, and maintained as an in-house breeding colony, originally provided by Dr. John Kopchick, Ohio University (32)) were injected via the lateral tail vein with AAV8-Null or AAV8-STAT5_CA_ (1.5 x 10^11^ GC per mouse, diluted in sterile PBS). Seven days later the mice were decapitated, and livers were collected. Nuclear protein was extracted using NE-PER Nuclear Cytoplasmic Extraction Reagent (cat. # 78833, Pierce, Rockford, IL). Western blots were performed to detect the Flag-tag sequence using rabbit monoclonal anti-DYKDDDDK Tag antibody (Cell Signaling #14793). To test the efficacy of AAV8-STAT5_CA_ to increase hepatic expression and restore circulating levels of IGF1 to normal levels, we used a mouse model with adult-onset hepatocyte-specific GH receptor knockdown (aHepGHRkd) (33), prepared as follows. Male GHR^fl/fl^ mice (10-12 wk old) were injected via lateral tail vein with 1.5 x 10^11^ GC/mouse of either AAV8-Null, to obtain GH receptor-intact control mice, or AAV8 expressing Cre recombinase (AAV8-TBGp-Cre, Penn Vector Core), to generate aHepGHRkd mice. The above AAV8 injections done either alone or in combination with AAV8-STAT5_CA_ (at 0.36 or 0.75 x 10^11^ GC), with the dose of AAV8-Null adjusted to equalize the total GC of AAV8 per mouse across groups. Fourteen days later, blood and liver were collected to assay circulating IGF1 levels (Mouse/Rat IGF-1 ELISA, 22-IG1MS-E01, ALPCO) and liver IGF1 mRNA levels by qPCR (33). In other studies, GH receptor-intact male mice (C57Bl/6 strain from Jackson Labs, 10 wk old) were treated with AAV8-Null or AAV8-STAT5_CA_ (0.75 x 10^11^ GC per mouse, iv) and 14 d later livers were collected and used for RNA-seq analysis (see below).

### AAV8-STAT5_CA_ dose response studies

Studies to test the ability of AAV-STAT5_CA_ to feminize hepatic gene expression were performed, in compliance with procedures approved by the Boston University Institutional Animal Care and Use Committee (protocol # PROTO201800698), and in compliance with ARRIVE 2.0 Essential 10 guidelines (34), including study design, sample size, randomization, experimental animals and procedures, and statistical methods. Male and female CD1 mice (Crl:CD(ICR), strain code 022), 7-8 wk of age, were purchased from Charles River Laboratories. Mice were housed in a temperature and humidity-controlled environment with a 12-h light, 12-h dark cycle, fed standard rodent chow, and supplied with tap water. AAV8-STAT5_CA_ or AAV8-Luciferase (control) (both produced in-house using methods described below) were administered to male mice by tail vein injection at doses ranging from 0.125 x 10^11^ to 2 x 10^11^ GC per mouse. Virus was diluted into PBS + 35 mM NaCl + 5% glycerol (stabilizer) to give a total injection volume of 150 uL. Mice were euthanized by cervical dislocation at 11 AM (lights on 7:30 AM-7:30 PM) to minimize the impact of circadian variations of gene expression (35,36).

### AAV8 production

AAV coding STAT5b_CA_ and Luciferase (control) was prepared and quantified by triple transfection of HEK293FT cells using methods adapted from protocols provided by Addgene (www.addgene.org). HEK293FT is a fast growing and highly transfectable derivative (cat. #R70007, ThermoFisher Scientific) of the SV40 large T-antigen-expressing HEK293T cell line commonly used for AAV production. On Day 0, 80% confluent flasks of HEK293FT cells were split 1:4 into DMEM culture medium [10% Fetal Bovine Serum (cat. #10437-028, Gibco), 1% PenStrep (cat. #15140-122, Gibco)] and then grown overnight to achieve 50-60% confluence at the time of transfection on Day 1. T182.5 flasks were each transfected with a total of 80 μg of plasmid DNA in a 1:1:1 (molar ratio) of three plasmids: pAAV-Helper helper plasmid, which encodes adenovirus genes E2A, E4 and (Cell Biolabs, VA); pAAV2/8 Rep/Cap plasmid (Addgene, plasmid #112864; replication protein derived from AAV-2 serotype, and Capsid protein from AAV-8); and either pENN.AAV.TBG.PI.N-FLAG-tag mSTAT5bCA.WPRE.bGH (see above) or pENN.AAV.TBG.PI.ffLuciferase.RBG (firefly luciferase expressed from TBG promoter; Addgene # 105538). Plasmids were mixed in a 3:1 weight ratio of total plasmid DNA:PEI in 4 mL of either OptiMeM™ (Gibco cat. # 31985-062) or serum-free DMEM per flask and then shaken vigorously for 30 sec. The mixture was incubated for 15 min at room temperature in a tissue culture hood, after which 4 ml of the mixture was added to each flask of ~85% confluent HEK293FT cells. After 24 h, the culture medium was replaced by DMEM containing 2% FBS and 1% PenStrep, and the cells were returned to the CO2 incubator for 5 days.

Virus was harvested on day 6 after transfection. A cell scraper was used to detach the cells from the flask, followed by collection of the media with cells in sterile 50 mL conical tubes. The tubes were centrifuged at 1000 x g for 10 min at 4°C, and the supernatant was decanted into a sterile bottle. Pellets were pooled and transferred to smaller tubes with 1x PBS + 200 mM NaCl + 0.001% Pluronic F68 (Poloxamer 188 Solution, cat. #P5556-100ML, Sigma) and pelleted again. Cell pellets were frozen at −80 °C for future use in purifying virus. The supernatant (which contains the majority of virus) was filtered through a 0.2 μM PES filter (VWR, cat. #97066-214). 25 mL of 40% PEG8000 (JT Baker, cat. #JTU222-09) was then added for every 100 mL of media supernatant, followed by overnight stirring at 4°C. The next day, the supernatant was incubated for 1 h at 4°C without stirring, to allow for full precipitation of virus before spinning at 2,818 x g for 15 min at 4°C. The supernatant was discarded and the PEG pellet, containing concentrated virus, was resuspended in 10 mL of 1x PBS + 0.001% Pluronic F68 + 200 mM NaCl by pipetting back and forth until the pellet was completely in solution.

The following buffers were prepared for iodixanol purification of the PEG-pelleted virus: (1) 1 M NaCl/PBS-MK Buffer (5.84 g NaCl, 26.3 mg MgCl2 and 14.91 mg KCl dissolved in 1x PBS to give a final volume of 100 mL), sterile filtered through a 0.2 μm PES filter; and (2) 1x PBS-MK Buffer (26.3 mg MgCl2 and 14.91 mg KCl dissolved in 1x PBS to a final volume of 100 mL and then filter-sterilized by passage through a 0.2 μm filter. Iodixanol gradients were prepared as follows: 15% Layer (4.5 mL of 60% Iodixanol (OptiPrep Density Gradient Medium, cat. # D1556-250ML, Sigma) + 13.5 mL of 1M NaCl/PBS-MK Buffer); 25% Layer (5 mL of 60% Iodixanol, 7 mL of 1x PBS-MK Buffer + 30 μL of Phenol Red (cat. #P0290-100ML, Sigma); 40% Layer (6.7 mL of Iodixanol + 3.3 mL of 1x PBS-MK Buffer; 60% layer (10 mL of 60% Iodixanol + 45 μL phenol red). A 10 mL syringe with an 18 g needle and a glass Pasteur pipet were used to layer the Iodixanol gradient in QuickSeal™ tubes (cat. # 344326, Beckman Coulter), from the bottom to the top, starting with 8 mL of the 15% layer, then 6 mL of the 25% layer, 5 mL of the 40% layer and 5 mL of the 60% layer (two tubes per AAV preparation from 10-12 T182.5 flasks). Resuspended virus-PEG pellet solution (5 mL) was added to the top of each gradient and 1x PBS was used to top off each tube before heat sealing. The tubes were spun for 90 min at 350,000 x g in a T70i Rotor at 10°C. The tubes were removed carefully (without disturbing the gradient) and secured to a clamp stand using a clamp. Twenty 1.5 mL Eppendorf tubes were opened and placed in a tube rack beneath the gradient in a biosafety cabinet. The QuickSeal™ tube was punctured at the 40-60% Iodixanol interface with an 18 g needle, using the bevel pointing up, and then the top of the tube was punctured with a 16 g needle. Factions of 1 mL were collected until the 25-45% interface was reached. 5 μL was removed from each fraction for analysis by qPCR to locate AAV-rich fractions before storing the remainder of each fraction at 4°C overnight. Each 5 μL fraction sample was treated with DNase I (cat #M6101, Promega) for 1 h at 37°C and then diluted 1:20 in 50 μg/mL yeast tRNA (cat #Am7119, Invitrogen). A known concentration of plasmid DNA used to prepare the virus was also diluted 1:200 in 50 μg/mL yeast tRNA and processed in parallel. AAV-containing fractions were pooled, typically giving a volume of 4-10 ml.

To concentrate the pooled fractions via buffer exchange, 12 mL of 0.1% Pluronic F68 in 1x PBS was added to a new Amicon Ultra-15 100K Centrifugal Filter Unit (cat. #UFC910024, EMD Millipore) and allowed to incubate at room temperature for 10 min. The Pluronic F68 in PBS was removed from the filtration unit and 12 mL of 0.01% Pluronic F68 in 1x PBS was added to the filtration unit before spinning at 3,000 RPM for 5 min at 4°C. The flow through was discarded and 12 mL of 0.001% Pluronic F68 in 1x PBS + 200 mM NaCl was added and spun at 3000 RPM for 5 min at 4 °C. The flow-through was discarded and up to 12 mL of sample was added to the column and spun at 3500 RPM for 8 min at 4°C. The flow through was discarded and more sample was added up to 12 mL, mixed well with the concentrated remaining sample and spun again. This continued until the sample was concentrated to a total final volume of 0.5-1 mL. The sample was washed with 10 mL of 0.001% Pluronic F68 in 1x PBS + 200 mM NaCl, using a pipette to mix thoroughly, and then spun again. The sample was concentrated to about 0.5 mL, collected using a pipette, and the walls washed with a small volume of 0.001% Pluronic F68 in 1x PBS + 200 mM NaCl to ensure all virus was collected. The concentrated virus was stored at 4°C for up to 2 wk, or at −80°C for long term storage.

Using 5 μL of the concentrated virus, purified AAV was treated with DNase I for 30-60 min at 37°C to eliminate any contaminating plasmid DNA remaining from the viral purification process. The DNase-treated viral samples were serially diluted 1:20 a total of five times in dilution buffer (50 μg/mL Yeast tRNA, 0.05% Pluronic F68) and the reference plasmid (used to prepare the virus) was serially diluted 1:10 a total of five times in dilution buffer. qPCR was performed as indicated above, using each plasmid dilution series to construct a calibration curve used to determine the titer of the virus.

### Liver RNA extraction and qPCR analysis

Total liver RNA was extracted from frozen mouse liver with TRIzol reagent (Invitrogen Life Technologies Inc., Carlsbad, CA), and nuclear RNA was purified from nuclei isolated from frozen liver tissue and then extracted with TRIzol LS as detailed elsewhere (37). RNA (1 μg) was treated with DNase I (cat. #M6101, Promega) to remove DNA contamination. cDNA was then synthesized using High-Capacity cDNA Reverse Transcription Kit (cat. #4368814, Applied Biosystems). qPCR was performed using primers specific to each RNA, designed using Primer Express or Primer3 software (http://bioinfo.ut.ee/primer3-0.4.0/), as follows: Sult3a1: 5’-CATTGTCACATATCCAAAGTCTGGT-3’ (forward), 5’-GAAAGGATCTGCTGGGTCCA-3’ (reverse) (oligonucleotides# 5735-5736); A1bg: 5’-GAACCCTCTGAGCCCAGTGA-3’ (forward), 5’-GAGTGGGTGGAGCCTGTGAG-3’ (reverse) (oligonucleotides# 5723-5724); Cux2: 5’-CCTCAAGACGAACACCGTCAT-3’ (forward), 5’-GCGCATCCTGGACCTGTAGT-3’ (reverse) (oligonucleotides# 1382-1383). Quantitative real-time PCR was carried out on a CFX384 Touch Real-Time PCR Detection System (Bio-Rad) using Power SYBR Green PCR Master Mix (ThermoFisher). Normalized linear Ct numbers based on 18S RNA as control were computed to determine the relative expression level of each gene across treatments.

### RNA-seq analysis

Polyadenylated mRNA was isolated from 1 μg of total liver RNA or from 1 μg of liver nuclear RNA from each treatment group (see below) using NEBNext Poly(A) mRNA Magnetic Isolation Module (cat. #E7490, New England Biolabs). The resulting polyA-selected RNA was used to prepare RNA-seq libraries using the NEBNext Ultra™ Directional RNA Library Prep Kit for Illumina (cat. #E7420L, New England Biolabs), NEBNext Multiplex Oligos for Illumina® Dual Index Primers (cat. #E7600, New England Biolabs), and AMPure XP Beads (cat. #A63881, Beckman Coulter Inc., Indianapolis, IN). RNA-seq libraries were prepared from two independent studies: 1) Sample series #G169: livers from 10-12 wk old male C57Bl/6 mice treated with 0.75 x 10^11^ GC of AAV8-STAT5_CA_ or AAV8-Null control then excised 2 wk later, with two sequencing libraries prepared for each treatment group, each derived from polyA-selected total liver RNA pooled from n = 2 or 3 individual livers (biological replicates); and 2) Sample series #G186: livers from 7-8 wk old male CD1 mice treated with 2 x 10^11^ GC of AAV8-STAT5_CA_ or control then excised 4-6 wk later, with a total of 9 sequencing libraries prepared, each one from an individual liver (n=5 AAV8-STAT5_CA_ libraries, n=4 control libraries) and derived from polyA-selected nuclear RNA prepared from nuclei extracted from frozen liver tissue, to increase the sensitivity for detection of the large fraction of lncRNAs that are nuclear-enriched and tightly bound to chromatin in mouse liver (37). Libraries were multiplexed and sequenced by Novogene, Inc (Sacramento, CA) to an average depth of 18-25 million 150 nt paired-end sequence reads on an Illumina HiSeq instrument. RNA-seq data was analyzed using a modified version of the custom pipeline described earlier (15). Briefly, sequence reads were aligned to mouse genome build mm9 (NCBI 37) using STAR aligner (38) and FeatureCounts (39) was used to count sequence reads mapping to the union of the exonic regions in all isoforms of a given gene (collapsed exon counting) based on an mm9 Gene Transfer Format file comprised of 75,798 mouse genes: n = 20,884 RefSeq protein coding genes, n=48,360 mouse liver-expressed lncRNAs genes (a much more complete listing than the set described previously in (40)), n=2,061 RefSeq non-coding genes (NR accession numbers) that do not overlap the set of 48,360 lncRNAs, n =4,490 Ensembl non-coding lncRNAs that do not overlap either the RefSeq NR gene set or the 48,360 lncRNA gene set, and n=3 other lncRNAs with interesting liver functions (lnc-LFAR1, LeXis, Lnclgr) (41) and in the gene annotations provided in **Table S1**. Raw sequencing files and processed data files are available at GEO (https://www.ncbi.nlm.nih.gov/geo/) under accession numbers GSE196014 and GSE196015.

### Differential expression analysis

EdgeR (42) was used to determine significant differential expression of RNA-seq data for the following comparisons: Males + AAV8-STAT5_CA_ at 2 x 10^11^ GC vs. Control males, and Males + AAV8-STAT5_CA_ at 0.75 x 10^11^ GC vs. AAV8-null males, with full datasets shown in **Tables S1-S3**. Further, a set of 475 sex-biased liver-expressed genes was defined as genes with male liver vs female liver differential expression significant at FDR < 0.01 (**Table S2**). A set of 8,246 liver-expressed genes whose differential expression is stringently sex-independent was defined based on |fold-change| for sex-difference < 1.2 and FDR > 0.1 (**Table S3**). All analyses reported here for sex-independent genes used this list of 8,246 genes. Finally, for female-biased genes, a percent feminization value was calculated based on each gene’s response to each of the following treatments: AAV8-STAT5_CA_ 2 x 10^11^ GC, AAV8-STAT5_CA_ 0.75 x 10^11^ GC, and continuous infusion of male mice with GH (cGH) for 7 or 14 d (20), as follows: % feminization = 100% [FPKM (treated male) – FPKM (control male)]/[FPKM (control female) – FPKM (control male)] (**Table S2**).

Raw RNA-seq data obtained from livers of hypophysectomized mice (15) (GEO accession number GSE66003) was re-analyzed to obtain expression data for the full set of 75,798 mouse genes, including 48,360 liver-expressed lncRNAs, described above. Two distinct classes of sex-biased genes were identified based on differential expression between livers of control and hypophysectomized male and female mice at FDR <0.05 (22): class I male-biased genes are genes that show significantly decreased expression in male liver after hypophysectomy due to their dependence on the male plasma GH pulsatile pattern for expression; and class II male-biased genes are genes that are repressed by the female pituitary profile and, consequently, they are significantly induced in female liver following hypophysectomy. We also identified corresponding sets of class I female-biased genes as those whose expression in female liver decreases significantly after hypophysectomy due to a requirement for the female, near continuous GH pattern for expression, and class II female-biased genes whose expression is repressed by the male pituitary profile and consequently show significant de-repression in hypophysectomized male liver (see Fig. 1 of (43)). The enrichment of STAT5_CA_-responsive genes in each set of hypophysectomy-responsive sex-biased genes was calculated compared to that of a background set of the corresponding class of STAT5_CA_-unresponsive sex-biased genes. Statistical significance was evaluated using Fisher exact test.

**Fig. 1.**
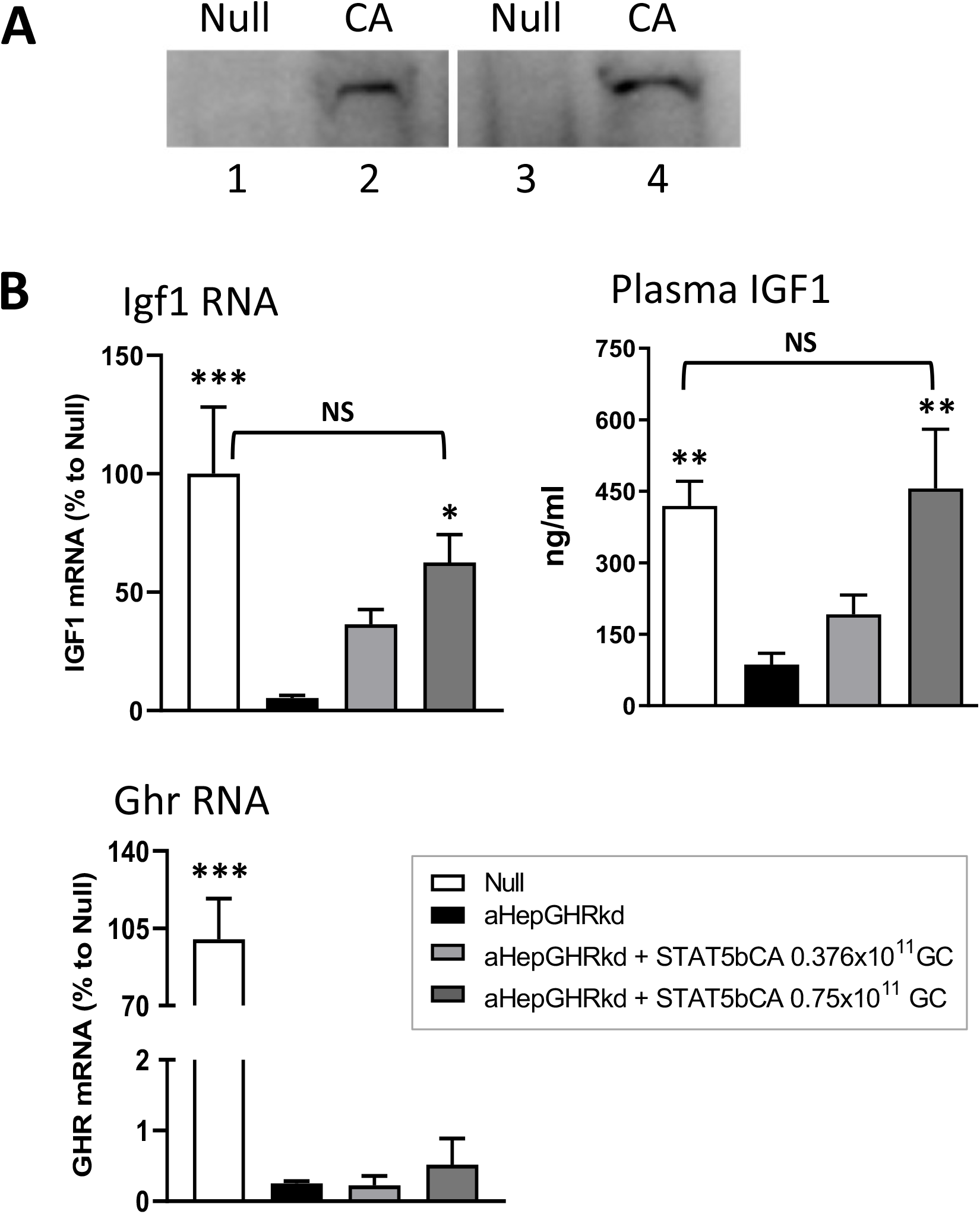
Functional validation of AAV8-STAT5_CA_ *in vivo*. (A) Nuclear localization of Flag-Tag STAT5_CA_ protein, detected by anti-Flag antibody on a Western blot of nuclear extracts from livers excised from mice 7 d after injection of AAV8-Null (control, lanes 1, 3) or AAV8-STAT5_CA_ (1.5 x 10^11^ GC, lanes 2, 4). Extracts were prepared from four individual livers. (B) Plasma IGF1 levels and liver Igf1 and GH receptor (Ghr) mRNA levels in male mice, GH receptor floxed control (Null) male mice, or male mice with adult-onset GH receptor knockdown (aHepGHRkd mice), with or without AAV8-STAT5_CA_ treatment at the doses indicated. Values shown are mean ± SEM for n =8 mice per group. Significance was evaluated by one way ANOVA with Bonferroni multiple comparison correction, implemented in GraphPad Prism. For comparisons to aHepGHRkd (second bar in each set): p< 0.001, ***; p< 0.01, **; and p=0.0595, *. NS, p > 0.05 for comparisons to Null, indicating restoration of Igf1 RNA and plasma protein.

### Enrichment of STAT5 ChIP-seq binding sites at STAT5_CA_-responsive genes

Male-enriched, female-enriched and sex-independent STAT5 binding sites (1,765, 1,790, and 11,531 sites, respectively) were obtained from our published ChIP-seq data for male and female mouse liver (9) (GEO accession number GSE31578). Each STAT5 binding site was mapped to the closest gene using the command bedtools -closest (44). All genes with a STAT5 binding site within 50 kb of the gene body were considered to be a STAT5 target. The enrichment of STAT5_CA_-responsive genes in each set of STAT5 binding site target genes was calculated by comparison to a background set of STAT5_CA_-unresponsive genes. Statistical significance was evaluated using Fisher exact test.

### Histology

Fresh liver tissue was submerged into 4% PFA and snap frozen. A portion of each liver was then placed in phosphate-buffered saline with 0.05% NaN3 for frozen tissue sectioning, and the remainder was transferred to 70% ethanol for paraffin embedding and H&E staining.

## Results

### Constitutively active STAT5b feminizes liver gene expression

GH-activated JAK2 catalyzes tyrosine phosphorylation linked to the activation and nuclear translocation of STAT5, which enables transcriptional activation of STAT5 target genes. GH-activated liver STAT5 is known to be essential for sex-biased liver gene expression; however, GH also activates other, STAT5-independent signaling pathways (24–26), which could contribute to the regulation of sex-biased gene expression. To determine whether the control of sex-biased liver gene expression by plasma GH pulses (male GH pattern) vs persistent GH stimulation (female GH pattern) is due to the downstream effects of pulsatile vs persistent STAT5 activation *per se*, we expressed a constitutively active form of STAT5 (STAT5_CA_) in male mouse liver to mimic the pattern of persistent STAT5 activation that occurs in female mouse liver (9). STAT5_CA_ cDNA expressed from the hepatocyte-specific thyroxin binding globulin promoter (45,46) was delivered using an AAV serotype 8 vector, which has high intrinsic tropism for the liver (30,31). We validated the AAV8-STAT5_CA_ viral vector by its ability to target STAT5 protein to the nucleus (**Fig. 1A**), which is a key feature of activated STAT5, and by its functional ability to increase hepatic *Igf1* RNA levels and circulating IGF1 in mice deficient in GH receptor signaling in hepatocytes (**Fig. 1B,** aHepGHRkd mouse liver).

Next, we investigated the impact of STAT5_CA_ on sex-biased gene expression in the liver. We delivered AAV8-STAT5_CA_ (2 x 10^11^ GC/mouse) or a control virus (AAV8-Luciferase) to young adult male mice and excised the livers 4-6 wk later. Liver gene expression was analyzed by nuclear RNA-seq, which enabled us to determine the effects of STAT5_CA_ on the nuclear transcriptome, including lncRNA genes, which are highly enriched in the nucleus owing to their tight binding to chromatin (37) (**Table S1**). AAV8-STAT5_CA_ substantially feminized the expression of liver sex-biased genes. Thus, 154 (59%) of 260 female-biased genes were induced and only 2 female-biased genes were repressed by AAV8-STAT5_CA_ in male liver. Furthermore, 114 (53%) of 215 male-biased genes were repressed and only 5 male-biased genes were induced by AAV8-STAT5_CA_ (**Fig. 2A**, **Table S2**). In contrast, only 1.1% of stringent sex-independent genes (**Table S3**) were induced and 0.7% were repressed by AAV8-STAT5_CA_. These trends, presented for the CD1 mouse liver model used extensively in our studies on liver sex differences (15,19,20,47), were confirmed in an independent study in C57BL6 mice, which were treated with AAV8-STAT5_CA_ at a lower dose and for a shorter duration (0.75 x 10^11^ GC for 2 wk) (**Fig. 2A,** bottom; **Table S2**). Feminization of the liver transcriptome by AAV8-STAT5_CA_ was dose-dependent, with partial feminization observed at virus doses as low as 0.125 x 10^11^ GC, as shown for three highly female-biased genes (**Fig. 2B**). Sex-biased genes responsive to AAV8-STAT5_CA_ were enriched for established sex-biased metabolic pathways, whereas the set of sex-independent genes responsive to AAV8-STAT5_CA_ was enriched for extracellular matrix/adhesion and glycoprotein/signal peptide genes (**Fig. 2C, Fig. 2D**).

**Fig. 2.**
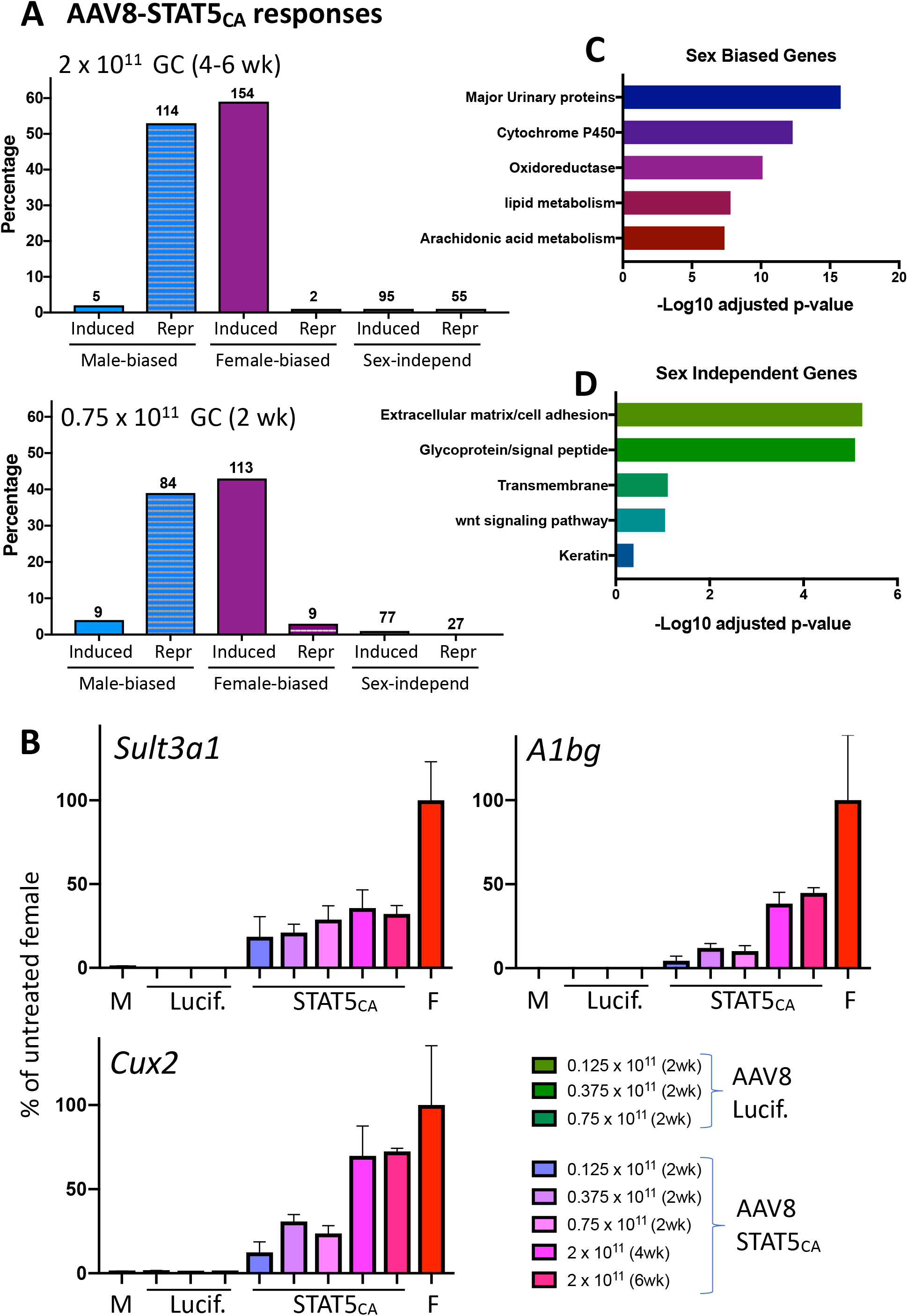
AAV8-STAT5_CA_ responsive genes in male liver. (A) Percentage of all genes that are either consistently sex biased at FDR < 0.05 (Table S2, n= 475 genes) or stringently sex-independent (SI) genes (Table S3, n = 8,246 genes) and whose expression in AAV8-STAT5_CA_-treated liver (as indicated) is significantly induced or repressed compared to control liver (FDR < 0.05). Top panel: results for CD-1 male mice; bottom panel: results for C57BL/6 mice. GC, genome copies of AAV8 virus. Number of responding genes is shown above each bar. (B) Dose responses for AAV8-STAT5_CA_ induction of three highly female-biased genes. Data based on RT-qPCR analysis of liver RNA, mean +/− SEM for n=2-6 livers per group, expressed as percentage of gene expression in control (untreated) female (F) mouse liver. M, male group. (C, D) DAVID pathway analysis of the sets of sex-biased genes (C) and sex-independent genes (D) that are responsive to STAT5_CA_. Shown are the top five enriched clusters, with bars indicating - log10 P values (Benjamini-Hochberg corrected) for enrichment.

### Role of STAT5 in liver feminization induced by continuous GH infusion (cGH)

cGH treatment overrides the endogenous male, pulsatile GH secretion pattern and progressively feminizes liver gene expression over a period of days (20). To understand the role of STAT5 in this process, we compared the changes in gene expression following AAV8-STAT5_CA_ treatment (2 x 10^11^ GC) to the changes seen in livers of male mice given cGH infusion for 7 or 14 days. AAV8-STAT5_CA_ and cGH infusion induced very similar changes in the expression of sex-biased genes in male liver (**Fig. 3A, Table S2**). Thus, 90% of the female-biased genes induced by AAV8-STAT5_CA_ were also induced by cGH (139/154 genes) and 81% of the male-biased genes repressed by AAV8-STAT5_CA_ were repressed by cGH (92/114 genes) (**Fig. 3B**). Overall, AAV8-STAT5_CA_ and cGH were both highly effective in feminizing liver gene expression, with median feminization values of 87% and 116%, respectively (**Fig. 3C, Table S2**). Female-biased genes that are highly induced (> 100-fold) by cGH infusion or by constitutive activation of STAT5 include the cytochrome P450 genes *Cyp3a16, Cyp3a41b, Cyp2c69*, and *Cyp2a4*, and other metabolic genes, such as *Fmo3* and *Hao2*. Of note, 62 female-biased genes were induced in male liver by cGH infusion but not by STAT5_CA_ and 62 male-biased genes were repressed in male liver by cGH infusion but not by STAT5_CA_ (**Fig. 3B, Table S2**). These genes may be regulated by GH signaling pathways independent of STAT5 (25). Finally, four male-biased genes that were repressed by cGH were unexpectedly induced by AAV8-STAT5_CA_ *(Chrna2, Fmn2, Nuggc, lnc5159;* **Fig. 3A**, **Table S2**).

**Fig. 3.**
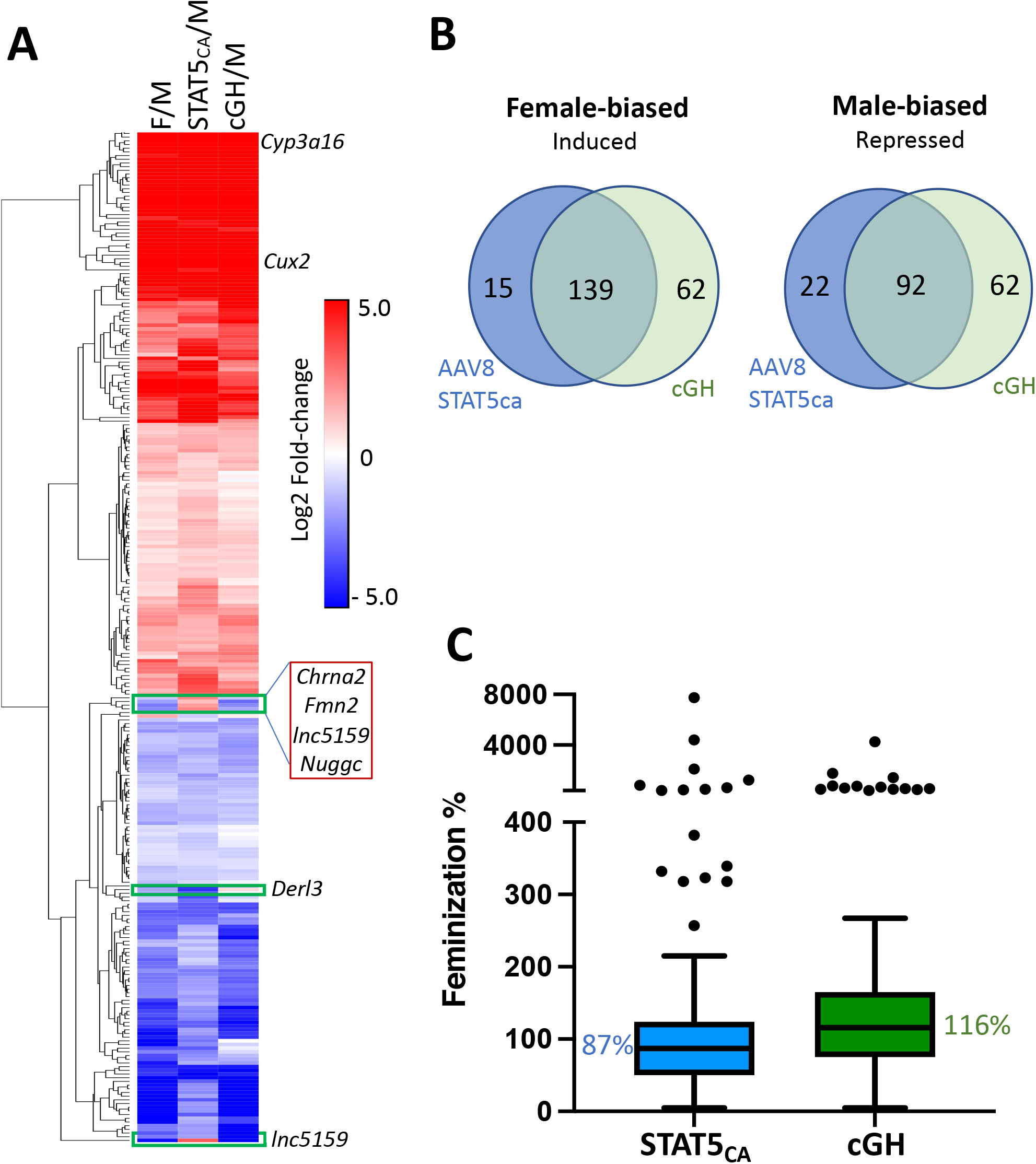
Effects of STAT5_CA_ and cGH infusion on sex-biased gene expression. (A) Heatmap showing impact of STAT5_CA_ or cGH infusion on the expression of 275 responsive sex-biased genes (log2 Fold-change increases (red) or decreases (blue)) in male mouse liver. M, male; F, female. (B) Overlap between female-biased genes induced by AAV8-STAT5_CA_ and those induced by cGH after either 7 or 14 days (left), and between male-biased genes repressed by AAV8-STAT5_CA_ and those repressed by cGH after either 7 or 14 days (right). Excluded are 5 male-biased genes induced by AAV8-STAT5_CA_ and 1 female-biased genes repressed by AAV8-STAT5_CA_. (C) Boxplots showing the distribution of feminization values for 139 female-biased genes induced in male liver in both mouse models (see Table S2). Median value, horizontal line in each box.

### STAT5_CA_ regulates many direct STAT5 gene targets

We identified an initial list of potential STAT5 target genes, defined as genes that met the following three criteria: 1) gene expression shows a significant change upon loss of STAT5, as seen in either male or female STAT5-deficient mouse liver (12,19); 2) gene expression is induced or repressed in either male or female liver following pituitary hormone ablation by hypophysectomy, which abolishes GH-dependent STAT5 activation (15); and 3) gene expression is restored when hypophysectomized mice are given an exogenous pulse of GH, which restores pulsatile STAT5 signaling in male liver (15). We filtered out genes showing an inconsistent response to GH treatment or STAT5 deficiency to obtain a final list of 227 putative STAT5 target genes, of which 115 showed sex-biased expression (**Table S4**). Importantly, 40% (90 of 227) of the predicted STAT5 target genes and 45% (52 of 115) of the sex-biased predicted STAT5 targets were responsive to AAV8-STAT5_CA_ (**Table S4**, column AC). These STAT5_CA_-responsive STAT5 targets both sex-biased genes and well-established sex-independent STAT5 gene targets, such as *Igf1, Socs2, Cish* and *Onecut1*, whose expression was increased in AAV8-STAT5_CA_-treated male mouse liver.

Next, we investigated whether responsiveness to AAV8-STAT5_CA_ is associated with local STAT5 binding to liver chromatin. We mapped liver STAT5 binding sites identified by chromatin immunoprecipitation (ChIP-seq) (9) to their putative target genes (nearest gene within 50 kb; **Table S2, Table S3**). These sites include many genomic regions where STAT5 binding to liver chromatin is significantly stronger in males than in females, and vice versa (male-enriched and female-enriched STAT5 binding sites, respectively) (9). For each gene class of interest, we determined the enrichment of STAT5 binding in the set of genes responsive to AAV8-STAT5_CA_ compared to the set of genes that were unresponsive to AAV8-STAT5_CA_. We observed a strong, significant enrichment of female-biased STAT5 binding at the set of female-biased genes induced by STAT5_CA_ as compared to the corresponding background set of female-biased genes not induced by STAT5_CA_ (**Fig. 4A**, **Table S5**). Male-biased genes repressed by STAT5_CA_ were depleted of STAT5 binding, but not significantly. The strong enrichment of female-biased STAT5 binding at female-biased genes induced by STAT5_CA_ indicates that it is the persistence of STAT5 binding, acting in a positive regulatory manner, that induces these female-biased genes in male liver. By contrast, we observed strong enrichment of sex-independent STAT5 binding at sex-independent genes that are repressed by STAT5_CA_, but not at sex-independent genes induced by STAT5_CA_ (**Fig. 4A**). Thus, in the genomic context of these sex-independent genes, STAT5 can act as a repressor, consistent with studies in GH-deficient rat liver (48). Finally, sex-independent genes induced by STAT5_CA_ were enriched for male-biased STAT5 binding, consistent with the positive regulatory potential of those sites evident from our earlier work (9).

**Fig. 4.**
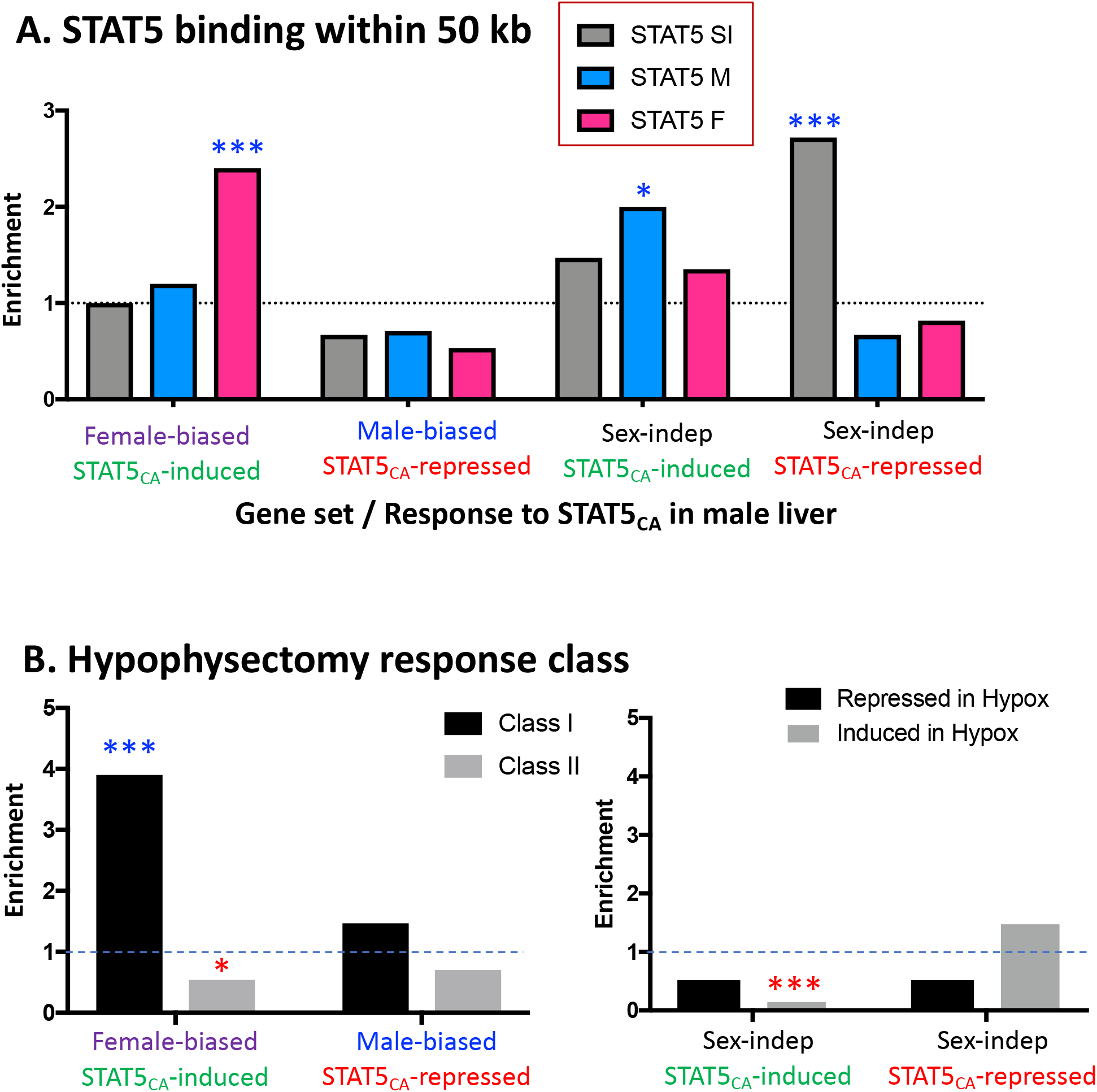
Enrichment of STAT5 binding sites (A) and hypophysectomy response class (B) for the sets of genes induced or repressed by STAT5_CA_. (A) Data are shown for STAT5 binding sites that are male-biased (M; 1,765 sites), female-biased (F; 1,790 sites), and sex-independent (SI; 11,531 sites), as defined by ChIP-seq, and mapped to their target genes (nearest gene within 50 kb). Enrichments are expressed as the ratio of STAT5_CA_-responsive genes compared to STAT5_CA_-unresponsive genes that are associated with each of the 3 sets of STAT5 binding sites. (B) Enrichment of class I or class II sex-biased genes (left) and sex-independent genes (right) in the STAT5_CA_-responsive gene sets, expressed as the ratio of the number of STAT5_CA_-responsive genes compared to the number of STAT5_CA_-unresponsive genes in each class (left) or that are induced or repressed by hypophysectomy (right). Significance was determined by Fishers exact test: *, p< 0.05, ***, P < 0.001. Blue asterisks, significant enrichment; red asterisks, significant depletion.

### STAT5_CA_ preferentially induces class I female-biased genes

In principle, STAT5_CA_ may act by either of two distinct mechanisms. By mimicking the actions of a persistent, female-like plasma GH profile, STAT5_CA_ may induce expression of female-biased genes that are positively regulated by the female GH pattern, i.e., class I female-biased genes. Alternatively, by eliminating the pulsatile nature of STAT5 signaling that is ongoing in male liver (15), STAT5_CA_ may effectively override the inhibitory activity that endogenous male plasma GH pulses exert on class II female-biased genes and thereby de-repress their expression (22). To distinguish these two models, we compared the hypophysectomy response class frequency of the female-biased genes that are induced by STAT5_CA_ to those that are not induced by STAT5_CA_ (**Table S2**, **Table S5**). We found that the set of STAT5_CA_-induced female-biased genes is significantly enriched in class I female-biased genes; furthermore, it is significantly depleted of class II female-biased genes (**Fig. 4B**, left). No significant hypophysectomy class enrichment was seen for STAT5_CA_-repressed male-biased genes. Sex-independent genes induced by STAT5_CA_ showed a highly significant 7-fold depletion of genes whose expression was induced when GH was ablated by hypophysectomy, whereas sex-independent genes repressed by STAT5_CA_ showed a tendency toward enrichment for genes induced by hypophysectomy (**Fig. 4B,** right), both consistent with the GH mimic nature of STAT5_CA_.

### Dose-dependent effects of AAV8-STAT5_CA_ on liver histology and *Igf1* expression

There were no consistent changes in hepatocyte morphology when AAV8-STAT5_CA_ was delivered at doses up to 0.75 x 10^11^ GC or in livers from mice given cGH infusion. However, when the dose of AAV8-STAT5_CA_ was increased to 2 x 10^11^ GC/mouse, we observed architectural distortion, hepatocyte hyperplasia, binuclear hepatocytes, karyomegaly and nuclear atypia (**Fig. 5**). To ascertain whether these responses are associated with STAT5 overexpression, we examined the expression levels of two classical STAT target genes in liver, *Igf1* and *Socs2*. Whereas AAV8-STAT5_CA_ induced the expression of *Igf1* moderately (2-2.5-fold increase above control liver levels at both doses), we observed strong induction of *Socs2*, which reached 8.5-fold at the lower dose and 14.4-fold above baseline at the higher dose (2 x 10^11^ GC), consistent with a functional overexpression of STAT5 activity at the higher AAV8-STAT5_CA_ dose (**Fig. 6**).

**Fig. 5.**
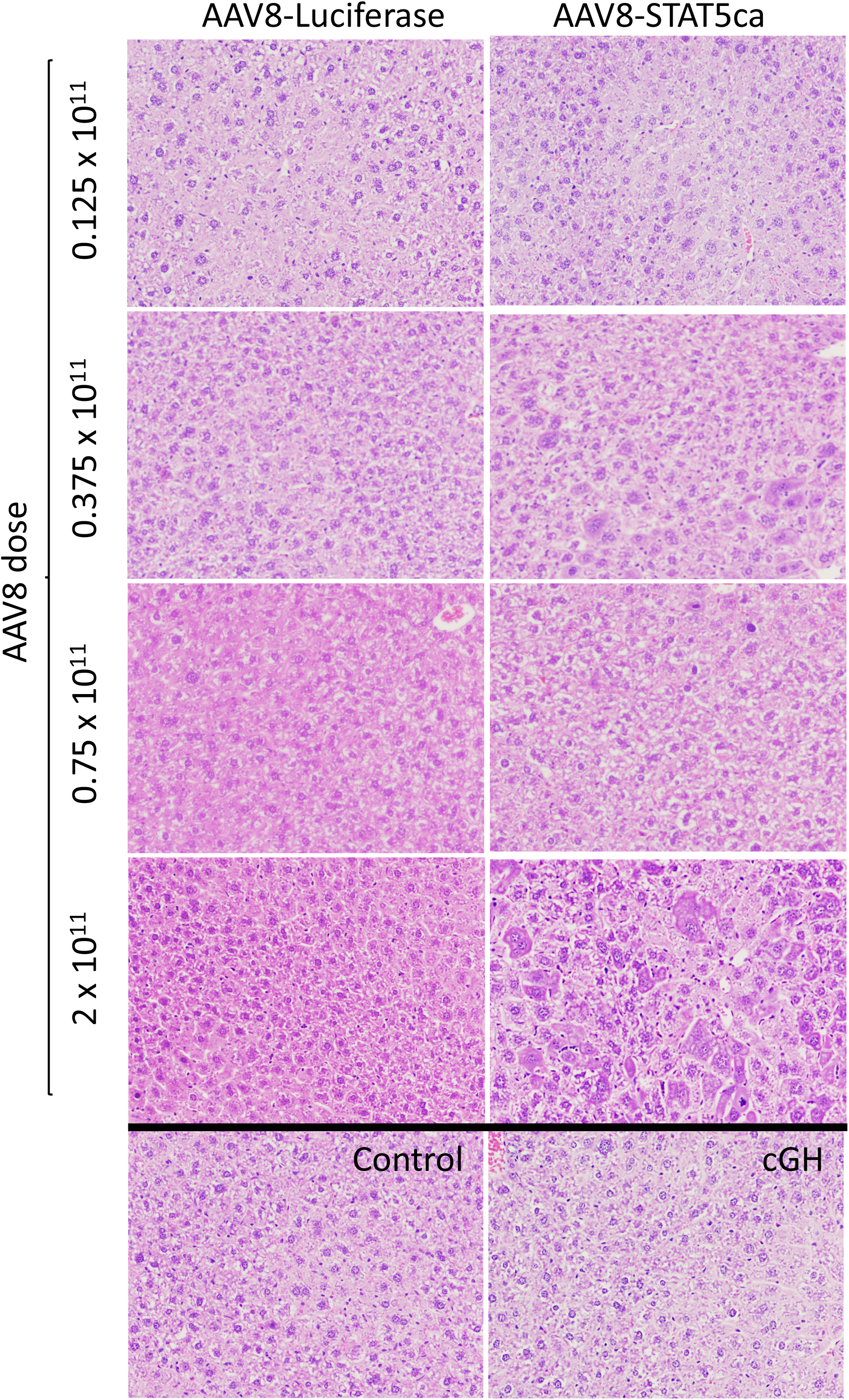
H&E staining of AAV8-STAT5_CA_-infected liver. Liver sections from mice treated with AAV8-STAT5_CA_ or AAV8-Luciferase at the indicated GC dose per mouse were stained with H&E, revealing a STAT5_CA_ dose-dependent histopathology, which was evident and most consistent at the highest dose of AAV8 dose. No histopathology was evident in mice given cGH infusion for 17 days. Images collected with an Olympus FSX-100 instrument at 16x.

**Fig. 6.**
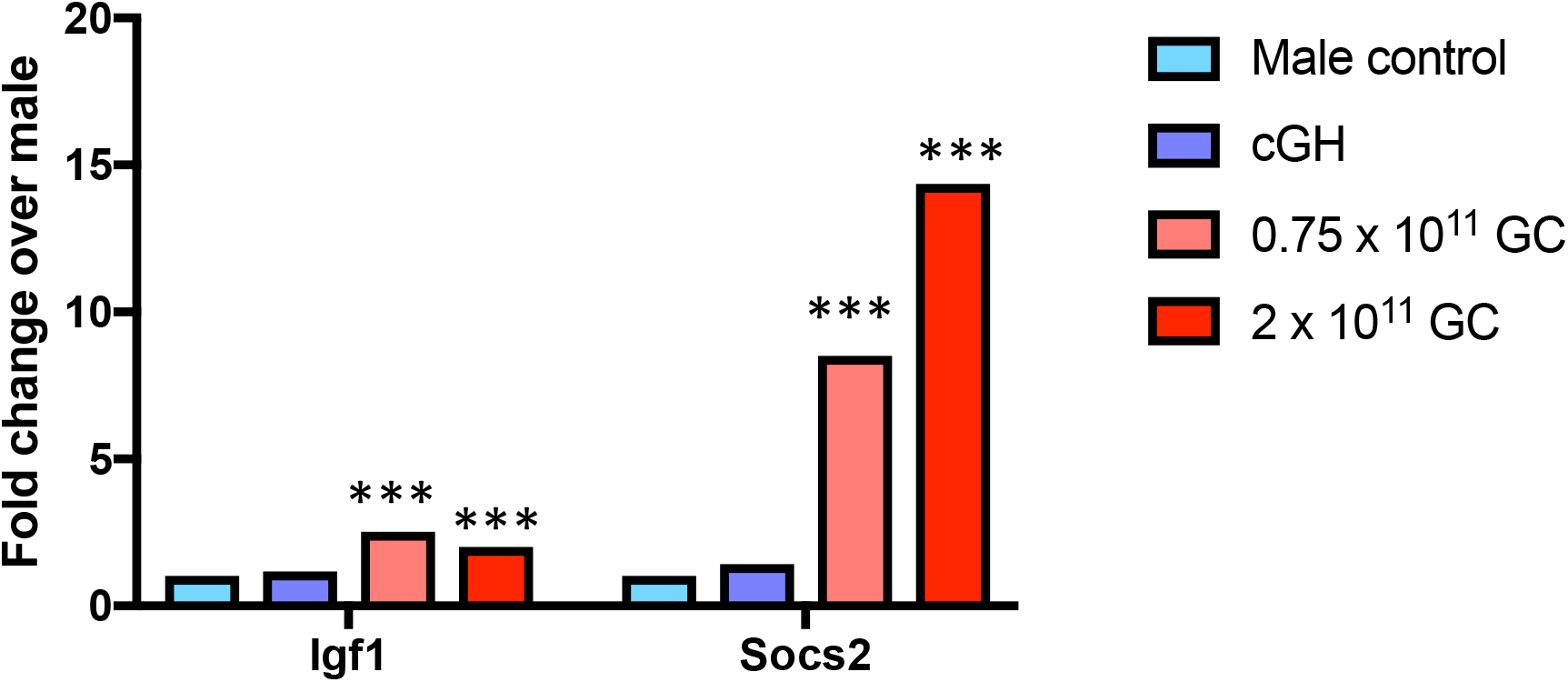
Expression of direct STAT5 target genes *Igf1* and *Socs2* in male liver: impact of cGH infusion and AAV8-STAT5_CA_. Shown are expression level determined by RNA-seq (fold-change values relative to male control), indicating supraphysiological induction of *Socs2*, and to a lesser extent *Igf1*, by AAV8-STAT5_CA_. Significance was determined by edgeR: ***, adjusted p-value < E-05.

## Discussion

The sex-dependent patterns of pituitary GH secretion – pulsatile GH release in males and persistent (near-continuous) GH release in females – activate hepatic GH receptor signaling to STAT5 in a manner that mirrors circulating GH profiles, i.e., pulsatile STAT5 activation and transcriptional activity in male liver, and persistent STAT5 activation in female liver. Here we show that persistent expression of activated STAT5, achieved using a liver-specific AAV8 viral vector to deliver the activated STAT5b mutant STAT5_CA_ to mouse hepatocytes *in vivo*, has extensive effects on sex-biased gene expression in the liver, with large numbers of male-biased genes repressed and female-biased genes induced. These effects are highly selective for sex-biased genes, insofar as fewer than 2% of genes showing sex-independent expression were impacted by STAT5_CA_ delivery. Many of the sex-independent gene responses are likely to be specific, in view of their enrichment for specific biological pathways related to extracellular matrix/cell adhesion and Wnt signaling.

Continuous infusion of GH in male mice (cGH treatment) mimics the endogenous female pattern of persistent circulating GH and overrides the endogenous pulsatile male secretion pattern, leading to substantial feminization of liver gene expression within 7 days (20,21). Here, we report a high overlap between gene responses to STAT5_CA_ expression and cGH infusion, providing strong evidence that persistent STAT5 activity alone is sufficient to account for a large fraction of the feminized genes. Nevertheless, some genes responded in a unique manner to either STAT5_CA_ expression or cGH infusion. While some of the differences in response to these two feminizing treatments may be due to cutoff thresholds, others clearly are not, for example *Cyp2b10* and *Cyp2c37*, whose expression in male liver was induced, and substantially feminized, by cGH infusion but which were unresponsive to STAT5_CA_ (**Table S2**). Differences between these two models for feminizing male liver gene expression may in part be due to STAT5-independent signaling pathways that are initiated by continuous GH, such as the GH receptor-dependent activation of PI-3 kinase, ERK signaling and Src family kinases (25).

STAT5_CA_ and cGH infusion both increased the expression of some female-biased genes to a level that exceeds levels of expression seen in untreated female liver. This phenomenon was somewhat more pronounced with cGH infusion (29 genes expressed at >2-fold the level of female liver after 14 days of cGH infusion vs. 18 genes exceeded that level following STAT5_CA_ treatment; **Table S2**). There are several possible reasons for the excessive feminization observed for these genes, and for the incomplete feminization observed for many other sex-biased genes. First, levels of STAT5 activity reached in AAV8-STAT5_CA_-infected hepatocytes may exceed the physiological levels of activated STAT5 in female liver, in particular at the higher dose of AAV8-STAT5_CA_ used in our study (c.f., overexpression of *Socs2;* **Fig. 6**). Related to this, both treatments used in this study, cGH infusion and expression of a persistently activated STAT5, differ somewhat from the physiological patterns in female liver, where there are short GH-off time periods, albeit not as prolonged or as frequent as the GH-off inter-pulse intervals that characterize males. Second, although STAT5_CA_ and cGH may both mimic female liver with respect to STAT5 activity and GH stimulation, they do not recapitulate the female hormonal environment with respect to other factors. One such factor is estrogen, which can also impact sex-biased gene expression in the liver, albeit it affects only 4% of liver sex-biased genes in a direct estrogen receptor-dependent manner (49). Indeed, it is remarkable how effective STAT5_CA_ and cGH are in feminizing male liver, despite the presence of a male gonadal steroid environment. Finally, intrinsic chromosomal sex differences associated with liver-expressed Y-chromosome-encoded epigenetic modifiers, such as *Uty* and *Kdm5d*, and partial X-inactivation/incomplete gene dosage compensation remain (50) and could also contribute to the observed differences between female liver and STAT5_CA_-treated or cGH-infused male liver.

Female-biased genes induced by STAT5_CA_ in male liver were significantly enriched for nearby female-biased STAT5 binding sites in liver chromatin when compared to the set of female-biased genes that were not induced by STAT5_CA_. Moreover, the STAT5_CA_-induced female-biased genes were significantly enriched for class I female-biased genes, whose expression in female liver is positively regulated by the female plasma GH pattern. Together, these findings lend strong support to the proposal that the de-repression of these female-biased genes in male liver involves direct binding of STAT5_CA_ to these STAT5-binding regulatory sites in association with induction of local, female-biased chromatin opening, which has been linked to female-biased STAT5 binding and gene activation (9,17). Female-biased genes not induced by STAT5_CA_ could be regulated by other mechanisms, including GH signaling independent of STAT5, as noted above. We did not observe a corresponding enrichment of STAT5 binding sites nearby male-biased genes repressed by STAT5_CA_, indicating that factors other than differences in STAT5 binding determine whether an individual male-biased gene will be repressed by STAT5_CA_ expression in male liver. Finally, sex-independent genes repressed by STAT5_CA_ showed strong enrichment for local sex-independent STAT5 binding sites, supporting the proposal that STAT5 can repress gene expression by a direct DNA binding mechanism.

We observed significant feminization of gene expression at AAV8-STAT5_CA_ doses as low as 0.125 x 10^11^ GC per mouse, i.e., 10-fold or more lower than standard doses of 2 x 10^11^ commonly used for AAV8-induced gene expression in mouse liver (31). This finding is in line with earlier studies in a GH-deficient rat model, where infusion of GH at a dose that results in as low as 3% of the nominal circulating female GH level restores the expression of certain female-biased genes, while other genes require higher GH doses and show only partial induction (51).

It is well established that a reduction of GH signaling is associated with liver steatosis and development of non-alcoholic steatohepatitis, both in humans and in mouse models, and increasing GH signaling can improve fatty liver endpoints (52,53). However, we found that treatment with a high dose of AAV8-STAT5_CA_ resulted in certain histopathological changes indicative of liver injury that cannot be attributed to the AAV8 infection *per se*, as they were not seen in AAV-Luciferase controls. Livers from mice given cGH infusion for 17 days did not show corresponding histopathological changes, despite the overall similarity of its effects on sex-biased gene expression. Conceivably, the histopathology associated with AAV8-STAT5_CA_ at the high dose could be related to the propensity of this viral vector to integrate into the host genome in the context of liver injury (54,55) or could be a supraphysiologic effect of STAT5 overexpression. Nonetheless, these observations highlight the importance of carefully evaluating such effects before considering STAT5_CA_ or other STAT5 derivatives for therapeutic use in treating liver disease.

## Acknowledgements

The authors thank the following Waxman lab members: Kritika Karri for providing annotations for the 48,360 mouse liver-expressed lncRNAs genes examined in this study, Dr. Maxim Pyatkov for RNA-seq analysis pipeline development, and Dr. Ravi Sonkar for assistance with histology imaging.

## Data availability

All data generated or analyzed during this study are included in this published article or in the associated Supplemental tables and the public data repository at GEO (https://www.ncbi.nlm.nih.gov/geo/) under accession numbers GSE196014 and GSE196015.

