## Supplementary material for "Constitutively active STAT5b feminizes mouse liver gene expression": Fig. S1 and Fig. S2

**Supplemental Figures:**

**Fig. S1.** UCSC genome browser screen shots showing STAT5 binding sites identified in male and female mouse liver (MH and FH, respectively) nearby class I female-biased genes induced by STAT5ca treatment of male liver (**A**), and sites nearby class I female-biased genes not induced by STAT5ca (**B**). Data are for male and female livers from mice collected at a peak of liver STAT5 activity [Y Zhang *et al*, Molec Cell Biol (2012) 32:880-896].

**Fig. S2**. H&E Staining of livers from three individual mice treated with AAV8-STAT5$_{CA}$ at 2 x $10^{11}$ GC/mouse and euthanized 4 weeks later. Shown and three fields at 4.2x and three fields at 16x for each liver. Histopathology is similar to the single liver shown in Fig. 5 for a mouse treated with AAV8-STAT5$_{CA}$ at 2 x $10^{11}$ GC/mouse and euthanized 6 weeks later. Images collected with an Olympus FSX-100 instrument.

**Supplemental Table:**

**Table S1**.  Impact of AAV8-STAT5ca on 42,938 liver expressed genes
**Table S2**. 475 Liver sex-biased genes
**Table S3**. 8,246 stringent sex-independent liver-expressed genes
**Table S4**. 227 putative STAT5 target genes
**Table S5**. Enrichment calculations supporting the data in Fig. 4A and Fig. 4B

Fig. S1

### A. STAT5ca-induced Female class I genes

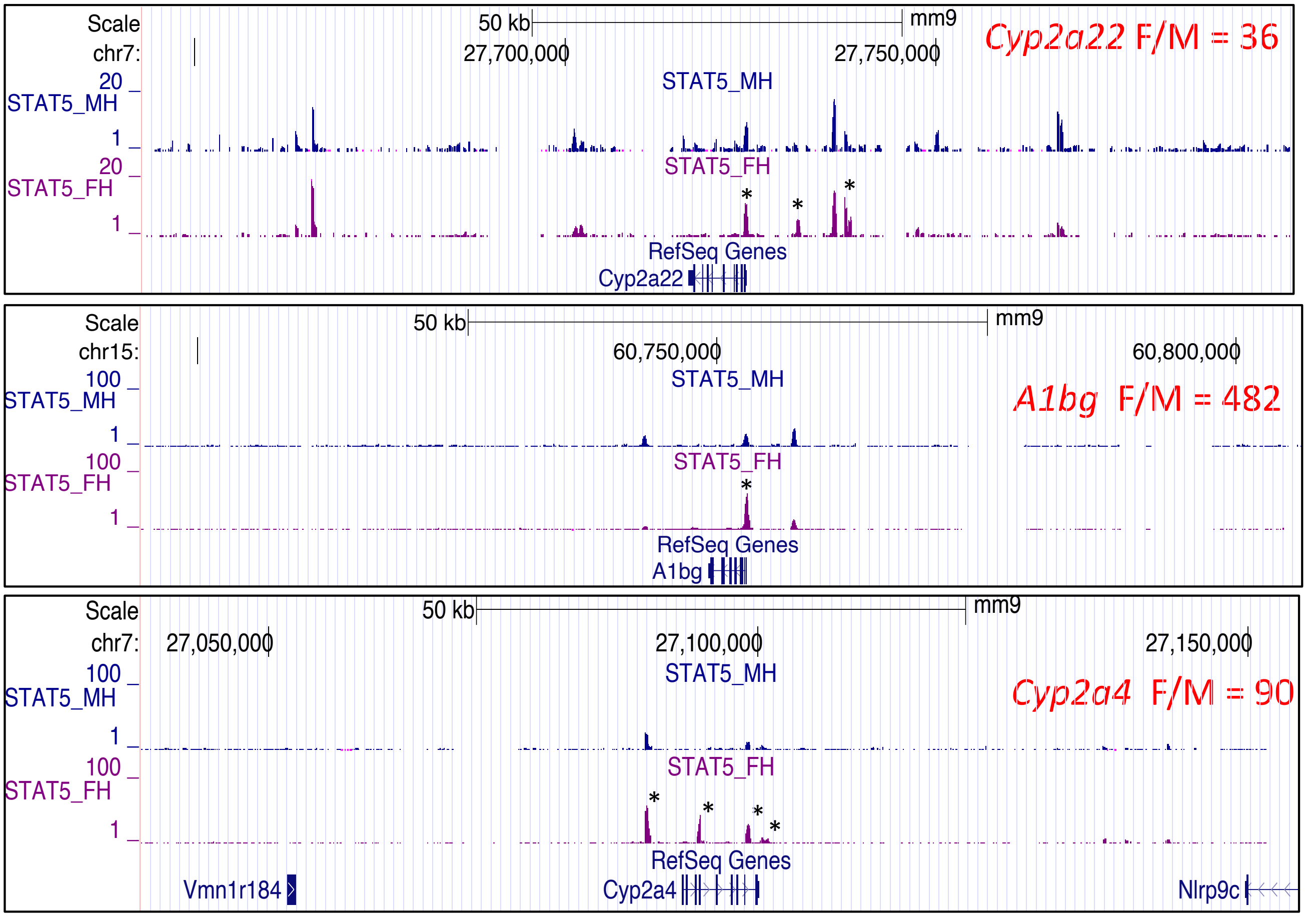


### B. STAT5ca-unresponsive Female class I genes

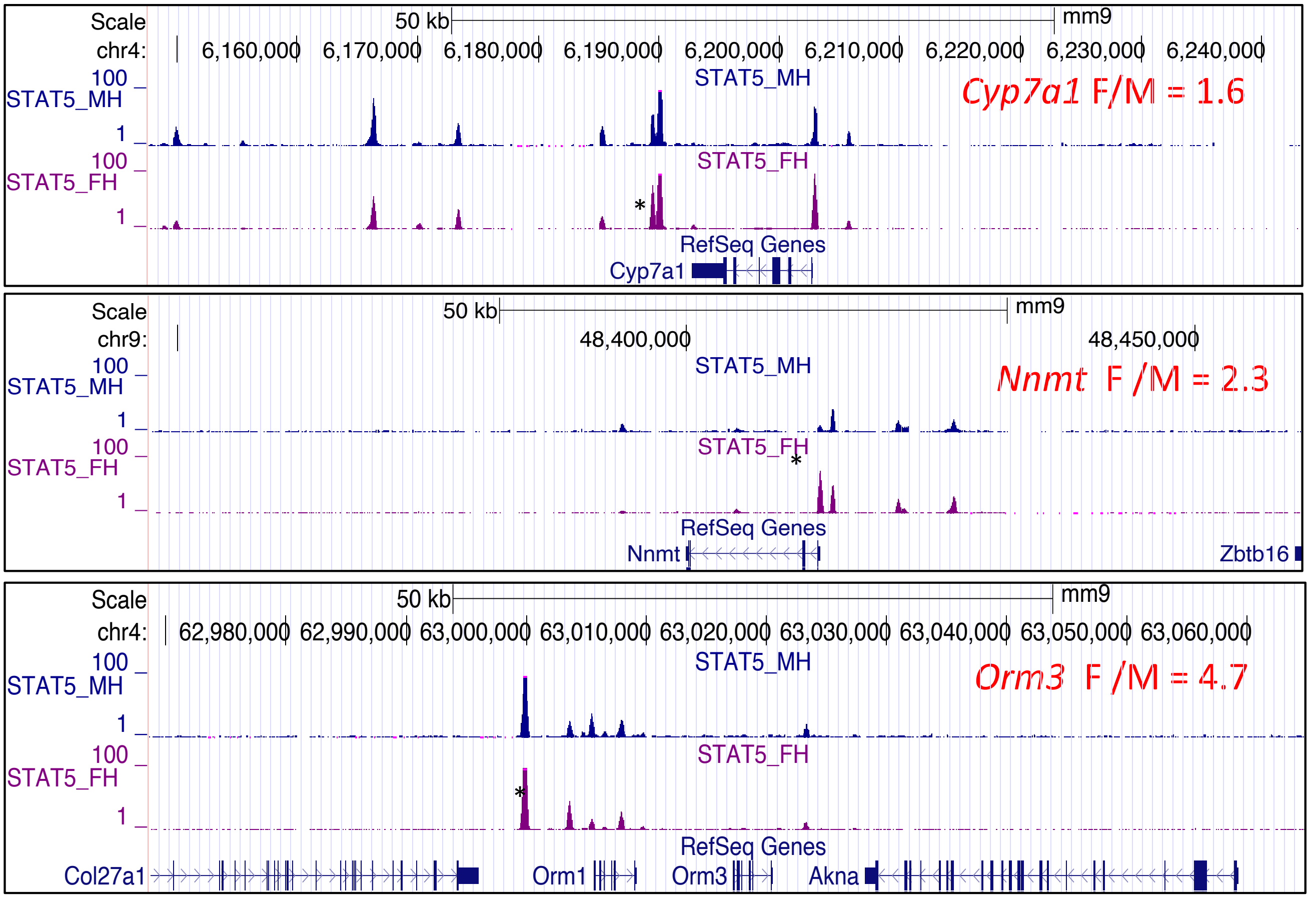

Fig. S2

**Liver 1:**

4.2X

16X

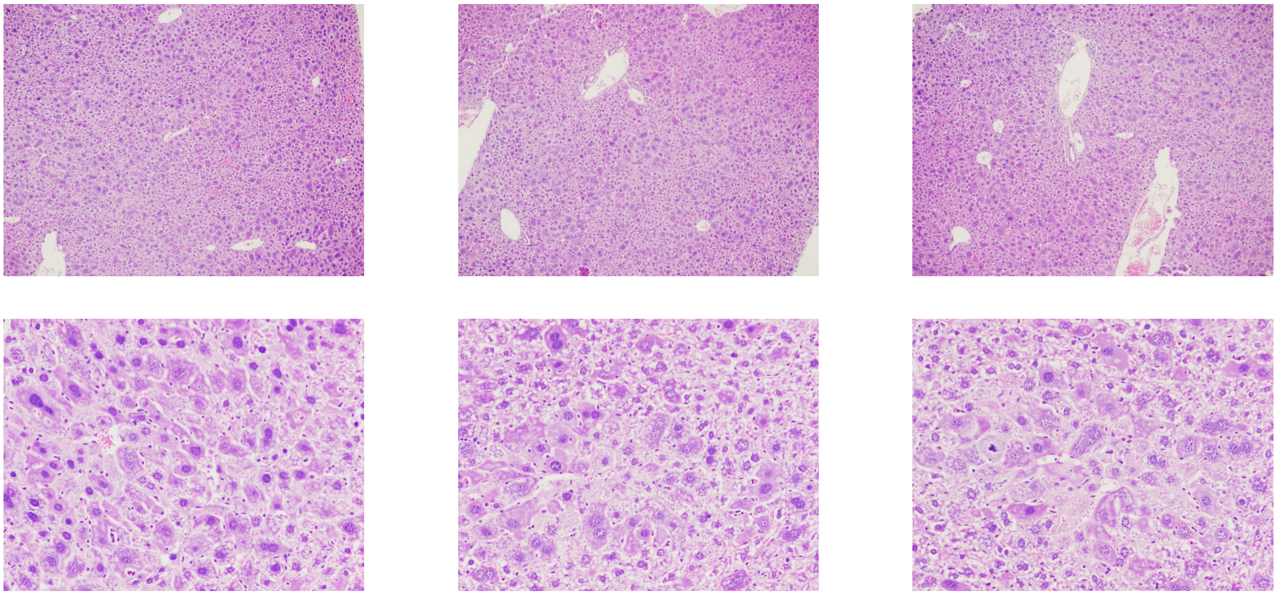

**Liver 2:**

4.2X

16X

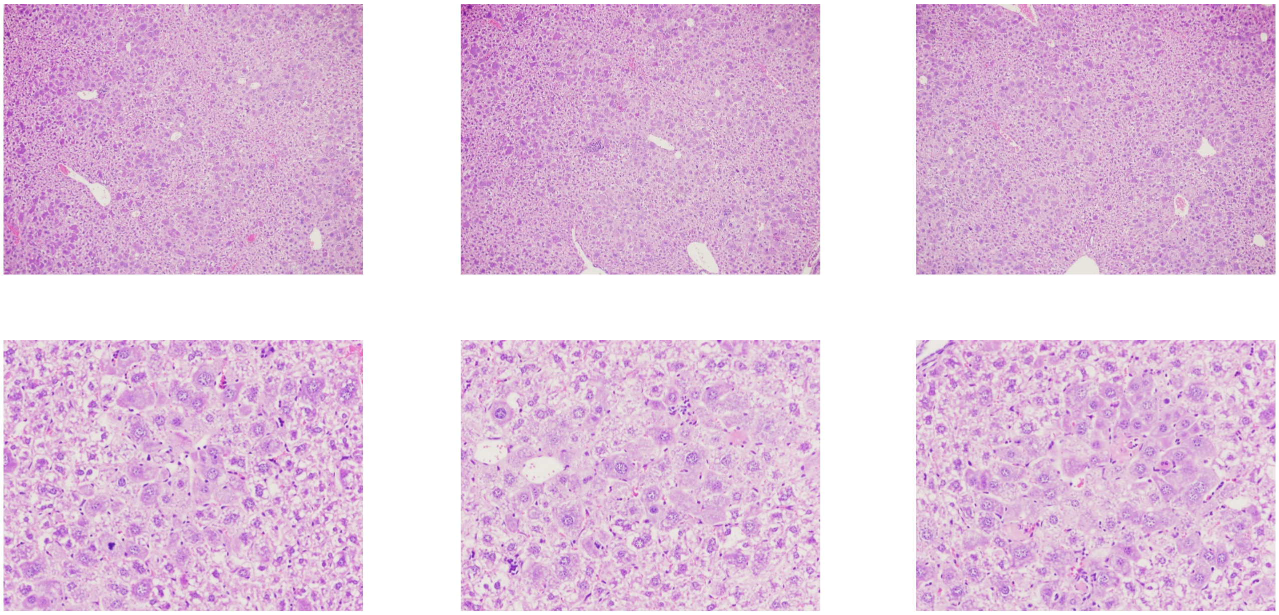

**Liver 3:**

4.2X

16X

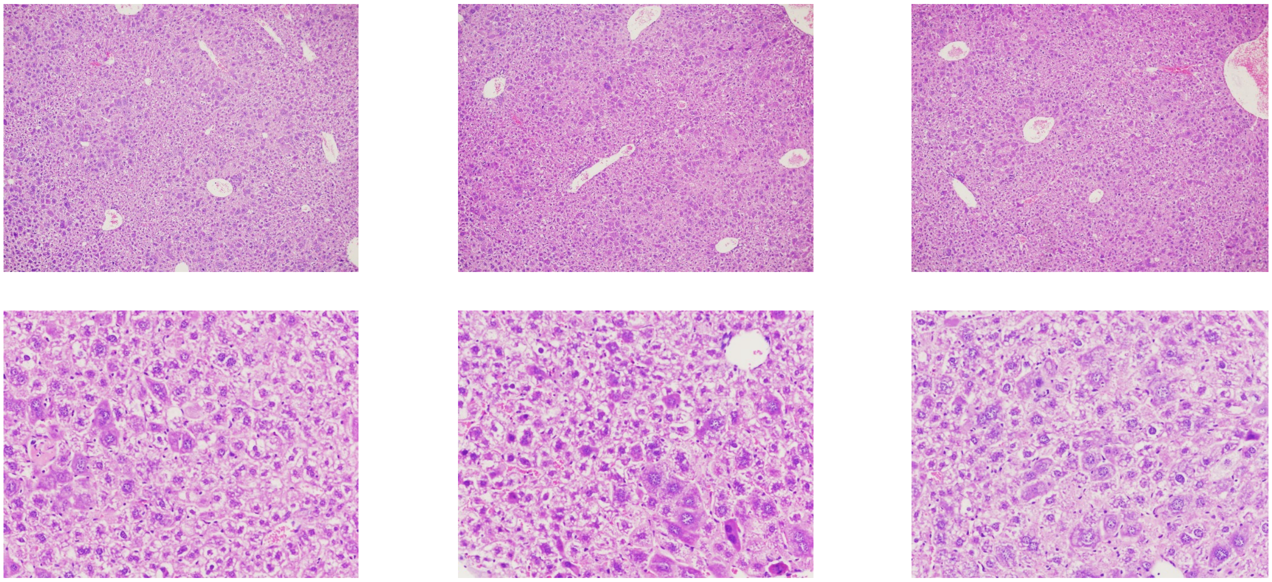
